# 3D-Printed Scaffolds for Local Chemotherapeutic Delivery in Resected Spine Metastases

**DOI:** 10.64898/2026.06.09.731191

**Authors:** Audrey A. Pitaru, Ateeque Siddique, Mansoureh Mohseni Garakani, Branda N. Boakye, Michael H. Weber, Abdellah Ajji, Michael Wertheimer, Isabelle Villemure, Lisbet Haglund, Derek H. Rosenzweig

## Abstract

Spinal metastases often occur secondary to breast, lung and prostate cancer and lead to instability, pain and poor quality of life. Standard care for spine metastases includes a multidisciplinary approach with surgery playing a major role in tumor resection, stabilization and decompression. Surgical resection with adjuvant is an effective treatment, yet it is often accompanied by tumor recurrence from residual disease. Furthermore, acrylic cements applied to defect sites provide stability, but they do not promote bone repair and can become destabilized during recurrence events. Developing new tools to stabilize defect sites, promote bone repair and locally deliver therapeutics may circumvent these limitations. We have previously developed mechanically competent 3D printed lactide/mineral scaffolds conducive to bone repair *in vivo*. We have also developed 3D printed nanoporous scaffolds conducive to both bone repair and chemotherapeutic delivery. Here, we set out to assess doxorubicin and cisplatin uptake and release rates and efficacy of drug delivery in 2D and custom physiological 3D cultures of two human cancer cell lines associated with metastases, MDA-MB-231 (human breast) and C42B (human prostate). Composite scaffolds had a compressive modulus close to trabecular bone, and could sustainably and effectively release doxorubicin and cisplatin as measured against both breast and prostate cell lines in 2D and 3D custom physiological metastases models. As a proof-of-concept, doxorubicin loaded composite scaffolds were implanted into rat caudal vertebrae following MDA-MB-231 xenograft resection. Following 6 weeks of implantation, no adverse events were noted and microCT analysis revealed boney integration of the construct. Taken together, these data indicate that our composite scaffolds may be an appropriate alternate therapy to stabilize bone defects, promote bone repair and effectively inhibit cancer recurrence post-tumor resection. Future work will test composite scaffolds using *in vivo* bone metastases models.

## Introduction

Cancer metastases remain a major challenge in patients with advanced cancer, and it is often the root cause of death, accounting for about 90% of cancer deaths [1]. Bone is a common site for cancer metastases, often arising from breast, prostate, and/or lung tumors [2]. Metastatic bone lesions often occur in the spine, where they are related to pain, instability and neurological deficits [3]. Surgical intervention is required for spine instability and spinal cord decompression, and the resulting bone defect is often large with poor healing. Acrylic cement is used to reconstruct defects, which can provide mechanical strength, but does not promote bone repair. Bone putty or ceramics can be used to reconstruct and repair smaller bone defects, yet they do not possess high mechanical strength. Moreover, local recurrence has been reported in over 30% of metastatic resection patients [4], indicating either incomplete resection or poor local uptake of systemic adjuvant chemotherapy. Improved surgical and medical care for spine metastases patients has resulted in increased survival times for these patients, which comes with increased risk of recurrence, further instability and the need for re-operation. There is therefore a need for multifunctional bone substitutes to stabilize defects, promote bone repair, directly deliver chemotherapeutics and block cancer recurrence.

Up to 34% of patients with surgically resected spine metastases experience site recurrence [5, 6]. This indicates that not all tumor cells are removed and that additional systemic medical oncology or radio-therapy treatments may not be effectively reaching the target site. Local therapeutic delivery devices may be appropriate for delivering high local doses of chemotherapeutics without the side effects associated with systemic doses [7, 8]. Materials such as polymers [9], lipids [10], polymeric hydrogels [11], and other types of scaffolds have all shown value in delivering chemotherapeutics such as doxorubicin for treating bone related cancers [12].Some of these materials are compatible with additive manufacturing, or 3D printing devices, providing a mechanism for generating scaffolds with appropriate mechanical and biochemical properties made to custom-fit in surgical resection sites.

3D printing has recently emerged as a cost-effective method to produce unique geometries combined with biomaterials to allow for patient-specific implants [13]. As for orthopedic applications, 3D printed scaffolds guide bone repair, can be fabricated using different materials and can locally deliver drugs [14]. A range of different biocompatible polymeric materials can be used to 3D print scaffolds, such as polymers, ceramics, and hydrogels. Biodegradable thermoplastic materials have been widely studied for 3D printing bone substitutes, yet they possess limitations as a singular material. Development of composite polymers has allowed for better mechanical properties and functionality that cannot be accomplished by single materials [15]. Poro Lay materials, such as Lay FOMM, are highly porous materials composed of a flexible thermoplastic polyurethane copolymer and polyvinyl alcohol (PVA). The PVA component is water-soluble, leaving a flexible, nanoporous, sponge-like structure suitable for *in vivo* use and drug uptake and delivery [16, 17]. Lactoprene is a commercial, 100% lactide medical-grade material similar to polylactic acid, a common filament used in 3D printing. Lactoprene is also available as a composite with (β-tricalcium phosphate) β-TCP, which can be used in clinical applications, and we have shown it to be suitable for *in vivo* repair [18].

In this study, we set out to develop a composite scaffold from two polymer types, a Lactoprene shell incorporating a Lay FOMM core, to support bone repair and locally deliver a high dose of chemotherapy. The Lactoprene provides a mechanically strong lactide-mineral material directing bone repair, and the sponge-like Lay FOMM core is loaded with therapeutics for local delivery. We tested scaffolds’ mechanical integrity to ensure that it has a comparable mechanical strength to trabecular bone, the chemotherapeutic release profile *in vitro* and the efficacy against breast and prostate cancer cell growth and viability. We hypothesized that 3D printed composite scaffolds would have comparable mechanical strength to trabecular bone and that the composite scaffolds with chemotherapeutics would be equally effective at inhibiting tumor cell proliferation and migration *in vitro* compared to direct chemotherapeutic treatment.

## Materials and Methods

### 3D Printing

3D-printed scaffolds were generated using a Monoprice MP Select Mini v2 (Monoprice, Inc; Brea, CA, USA). All scaffolds were designed using SOLIDWORKS 2015 (Dassault Systèmes, SolidWorks Corporation, Waltham, MA, USA), a Computer Aided Design (CAD) software. The height and diameter of the scaffold was set to 2 mm and 3 mm, respectively. The center hole was 2 mm, and the holes on the sides of the scaffolds were set to 0.5 mm, all in an orthogonal design (Figure 1a). A .stl file was generated and “sliced” to a G code using Ultimaker Cura 4.3.0 (Ultimaker B.V.; Utrecht, Netherlands). The filament used for 3D printing was Lactoprene-100M with βTCP (Polymed Inc), as described previously [18]. The printing parameters consisted of a 0.3 mm nozzle set to a temperature of 190°C, a bed temperature of 65°C, a printing speed of 20 mm/s, and the infill was set to 100%. The layer height was set to 0.175 mm. Lay FOMM 60 for the central pore was from MatterHackers (Burbank, CA, USA), which we have previously described [19, 20].

**Figure 1.**
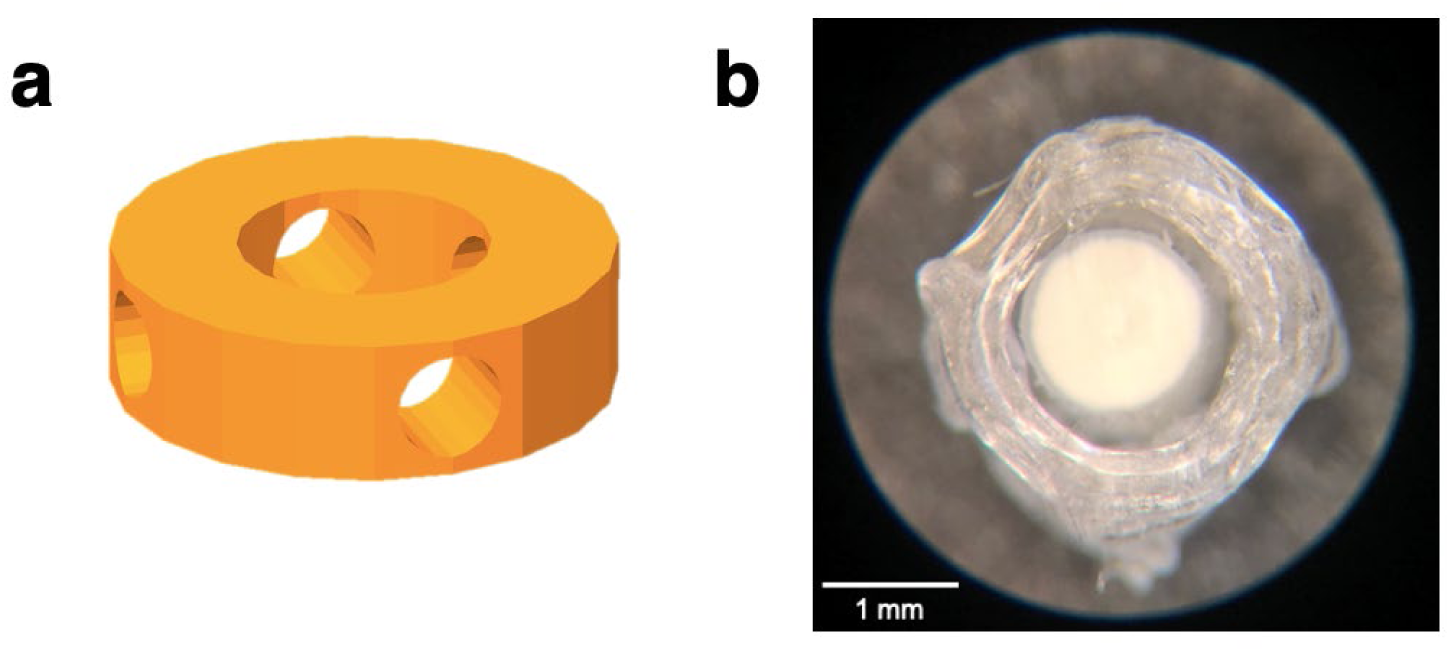
Scaffold visualization. (a) 3D CAD model design of outer scaffold shell. (b) Stereomicroscopy image of 3D printed scaffold made of Lactoprene shell with Lay FOMM 60 core insertion, viewed from top of scaffold. Scale bar = 1 mm.

To efficiently cut the Lay FOMM for placement in the center of lactoprene scaffolds, a 3D printed jig measuring tool was made with a 1 mm indent to ensure filament pieces inside the Lactoprene scaffold were uniform. Lay FOMM 60 filament was placed in the indented tool, and cut to 1.75 mm pieces using a razor and individual pieces were positioned inside the center of the Lactoprene scaffold with forceps. The composite scaffolds were placed in a 15 mL centrifuge tube with double distilled water for washing out PVA from Lay FOMM with two water changes per day over 3 days. Scaffolds were finally placed in 70% ethanol for 15 minutes for quick disinfection. Scaffolds were also placed under UV lighting for 15 minutes, turned over, and exposed for another 15 minutes and then stored in a sterile 15 mL conical tube. Sterilization steps are shown in Figure 2b. High-resolution imaging of the scaffolds was done using a Leica MS5 stereomicroscope with a 4x objective.

**Figure 2.**
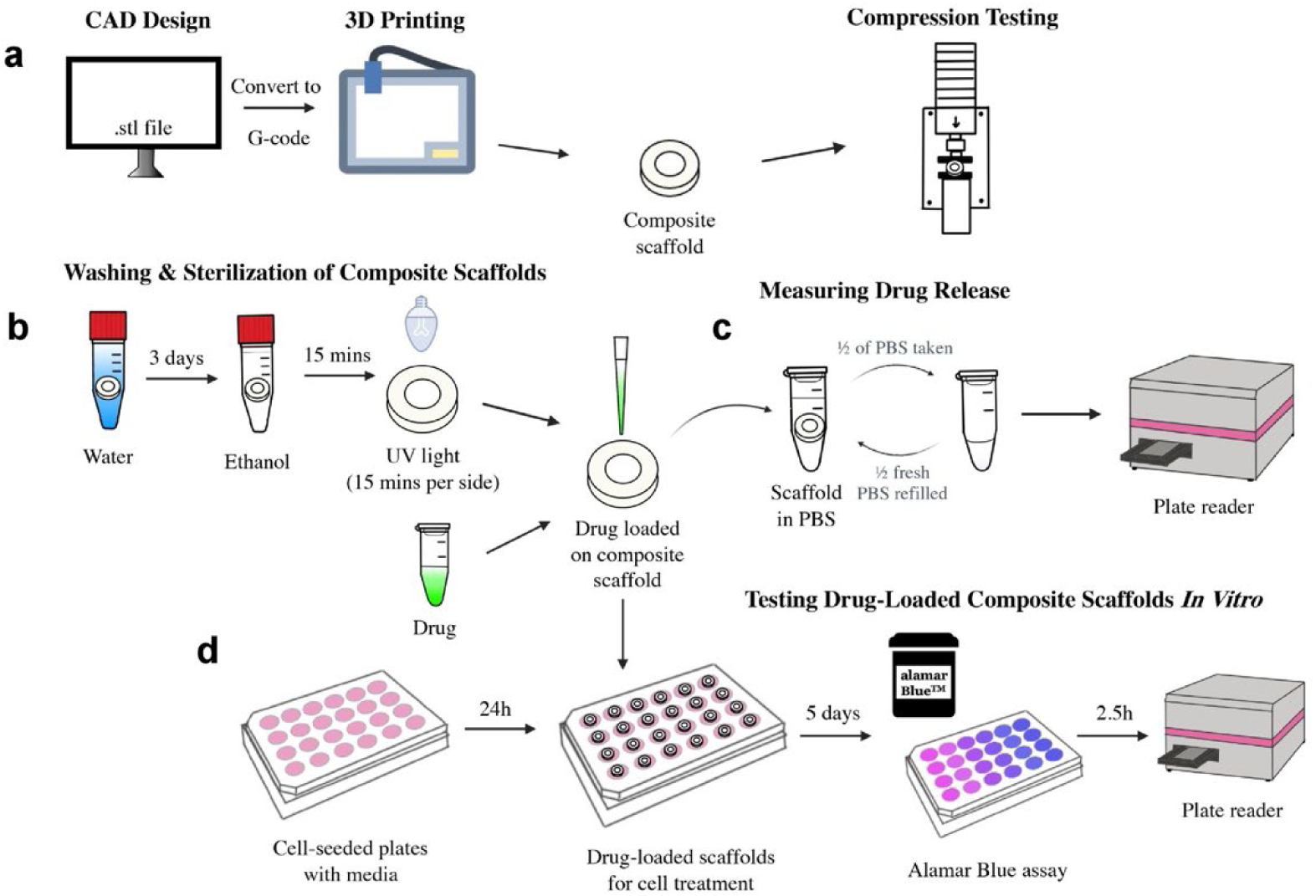
Schematic of (a) 3D printing and compression testing of scaffolds; (b) washing and sterilization of composite scaffolds; (c) drug release from scaffolds measured by fluorescence; (d) in vitro treatment of cancer cells with drug-loaded scaffolds and metabolic activity assessment via AlamarBlue assay.

### Compression Testing

Lactoprene-only (n=5) and Lactoprene + Lay FOMM 60 (n=9) scaffolds were fabricated and measured using a caliper to ensure scaffolds were placed under unconfined compression testing using a Mini-Bionix 858 (MTS; 14000 Technology Dr. Eden Prairie, MN) with TestStar II software as previously noted [18]. Briefly, the system had a 10 kN capacity, calibrated to 2 kN, with hysteresis at 0.15% and nonlinearity at 0.3% of full scale. Force and displacement data were recorded at 10 Hz. Tests used a strain rate of 1%/s and speed of 0.02 mm/s for 0.6 mm deformation (30% strain), starting at 2.1 mm and compressing to 1.4 mm. Young’s modulus was calculated from the linear region of stress-strain curves; yield stress and strain were determined using a 0.02% offset and end of the linear region, respectively as previously shown [21].

### Cell lines and Chemotherapeutics

One of the cell lines used in this study was the triple-negative MDA-MB-231 breast cancer cell line expressing luciferase (MDA-MB-231/Luc), provided by the laboratory of Joan Massagué at Memorial Sloan Kettering cancer center. As well, MDA-MB231 cells expressing green fluorescent protein (GFP; MDA-MB-231/GFP) and IMR-90 mCherry fibroblasts (red fluorescent protein; RFP), were previously reported [22–24]. The prostate cancer cell line C4-2B was from American Type Culture Collection (Manassas, VA, USA). All cells were cultured in DMEM with the exception of C4-2B cells which were cultured in RPMI cell medium, both from Thermo Fisher (Burlington, ON). Cells were cultured in complete cell medium containing 10% fetal bovine serum (FBS) and 1% penicillin-streptomycin (Thermo Fisher). For *in vitro* assays, low-serum cell medium was used, which contained 1% FBS. The chemotherapeutics used were doxorubicin hydrochloride (Sigma-Aldrich) and cisplatin (Selleckchem).

### Drug Release from Scaffolds

To evaluate the drug release profile of doxorubicin from the 3D printed scaffolds, three groups of scaffolds were prepared. Scaffolds were loaded with either PBS (control), 175 ng or 350 ng of doxorubicin. To load the drugs and PBS onto the scaffolds, the previously sterilized scaffolds were loaded with a 2 µL droplet of doxorubicin (containing either 175ng or 350ng Dox) or PBS that was placed directly on top of the Lay FOMM core which absorbed like a sponge and left in biosafety cabinet to dry for 45 minutes. 300 µL of PBS was placed in labled Eppendorf tubes and dried scaffolds were placed into the prepared tubes and stored in a 37°C incubator. Medium was taken from the tubes at 5, 24, 48, 72, 120, and 168 hours for measuring drug release. Each time solution was removed, equivalent volume of fresh PBS was replaced. Doxorubicin fluorescence intensity was measured using a TECAN Infinite M200 Pro microplate reader as previously described [19]. The concentrations of doxorubicin were interpolated from a standard curve. The measurement of drug release overview can be found in Figure 2c.

### Dose-Response

To assess the IC_50_ of Doxorubicin and cisplatin against the MDA-MB-231/GFP and C4-2B cancer cell lines, dose-response assays were conducted. 20,000 cells/well were plated in 48-well plates in complete culture medium. Medium was replaced with 300 µL fresh medium containing of various concentrations of doxorubicin or cisplatin in low-serum medium, as indicated in the figures ; PBS served as a vehicle control. After 48h, the metabolic activity was assessed via AlamarBlue (Invitrogen) resazurin reduction assay as previously described [22]. Briefly, 100µL of a 10% AlamarBlue solution prepared in low-serum medium was added, and cells were incubated for for 3h at 37°C. Fluorescence intensity of the metabolized AlamarBlue was measured using a TECAN Infinite M200 Pro microplate reader.

### In Vitro Treatment

The response of MDA-MB-231/Luc and C4-2B cell lines to drug-loaded scaffolds was determined by direct scaffold-cell incubation. 20,000 cells were plated in 48 well plates and, prepared scaffolds were added in low-serum medium. Four independent experiments were conducted for each cell line. Cells were treated separately with doxorubicin or cisplatin, using concentrations based on IC_50_ values from dose-response assays. Doxorubicin-loaded scaffolds for MDA-MB-231/Luc cells were prepared with 2 µL of 150 µM of doxorubicin; the direct positive control treatment of doxorubicin was 1 µM of drug added directly to medium. Since cisplatin could not be measure by fluorescence like doxorubing, cisplatin was added to scaffolds in varying concentration in 2 µL of medium (Table 1). 32 µM cisplatin was delivered directly in medium as a positive control. For C4-2B cells, doxorubicin-loaded scaffolds contained 2 µL of 30 µM dox, with a 0.2 µM directly treated control; cisplatin scaffolds used various concentrations (Table 2), with 8 µM as the directly treated control. Cells were incubated with scaffolds in 500 µL of low-serum medium per well. After 5 days, metabolic activity was measured via AlamarBlue assay.

**Table 1.** Scaffold cisplatin loading concentrations for in vitro MDA-MB-231/Luc cell treatment.

| Scaffold | Cisplatin Concentration Loaded ($\mu\text{M}$) |
| --- | --- |
| A | 2,667 |
| B | 5,333 |
| C | 8,000 |
| D | 10,666 |

**Table 2.** Scaffold cisplatin loading concentration for in vitro C4-2B cell treatment.

| Scaffold | Cisplatin Concentration Loaded ($\mu\text{M}$) |
| --- | --- |
| E | 1,200 |
| F | 1,867 |
| G | 2,667 |
| H | 5,333 |

### 3D In Vitro Treatment – Mixed Migration Model

To assess the migration of cells in a 3D model using 3D printed composite scaffolds as drug carrier a mixed migration model (MMM) was conducted. A previously developed 3D interface model (PP-3D-S) comprising a plasma-modified, electro-spun, nanofibrous 3D scaffold seeded with stromal cells and a 1% alginate-7% gelatin (A1G7) hydrogel embedded with tumor cells to mimic the cancer microenvironment as previously described [22–24]. Briefly, PLA (NatureWorks 4032D, density = 1.24 g/cc) was electrospun into nanofibers (600-800nm), cut into 9 mm disks, disinfected with RPMI medium + 1% antibiotics, and placed in non-adherent 48 well plates (SARSTEDT). Each well was seeded with 20,000 IMR-90 mCherry fibroblasts, incubated 30 min in 5% CO2, washed, topped with 100 µL of A1G7 containing 50,000 MDA-MB231/GFP tumor cells. Then, 200 µL of 100mM calcium dichloride (C7902, Sigma) was added for crosslinking, washed 10 minutes, and 500 µL of media was added per well.

The same steps were followed for the direct treatment (no 3D scaffold) . MMM groups included a control 3D printed scaffolds loaded with PBS, two differing doxorubicin doses (500 µM, 600 µM) on scaffolds, and direct doxorubicin treatments (1 µM, 2 µM), based on IC_50_values from this and previous studies [22–24]. Click or tap here to enter text.3D printed scaffolds were placed atop PP-3D-S scaffolds; after 5 days, the AlamarBlue assay was performed to assess metabolic activity, and drug-loaded scaffolds were discarded.

The migration/invasion of GFP-labeled tumor cells into the mCherry-fibroblast-seeded nanofibrous scaffold was assessed post-assay via fluorescence imaging (EVOS M5000, 4x objective, Thermo Fisher, Burlington, ON). Hydrogel was gently scraped, scaffolds fixed in 4% paraformaldehyde (PFA) for 15 minutes, washed with PBS, mounted on slides, and imaged. Migrated cells were quantified in 20 locations per sample using imageJ across three replicated experiments.

### In vivo testing

#### Tumor plug preparation

To prepare the composite hydrogel, sodium alginate (FMC BioPolymer, Protanal LF 10/60 FT) and Type B gelatin from bovine skin (Sigma-Aldrich, #G9391) were sterilized under UV light overnight, then dissolved in phosphate-buffered saline (PBS; Gibco, #10010). Solutions were stirred at 60 °C for 1 hour and then at room temperature for 2 hours, yielding a 3% w/v alginate and 7% w/v gelatin formulation (A3G7). The solution was centrifuged (1500 rpm, 5 minutes) to remove air bubbles and stored at 4 °C for up to one week. For tumor plug preparation, MDA-MB-231/Luc cells were suspended in 50 µL of supplemented DMEM and incorporated into 3 mL of thawed A3G7 to achieve a final cell density of 4 × 10⁶ cells/mL. The mixture was homogenized by syringe-coupling (BD, #309657; Ibidi, #10823) using 20-30 alternating transfers under sterile conditions. The A3G7-cell mixture was extruded through a 15G blunt needle into sterile 100 mM CaCl₂ (Sigma-Aldrich, #223506), crosslinked for 2 minutes, rinsed in PBS, and cut into 3 mm segments. The plugs were incubated overnight at 37 °C, 5% CO₂ in supplemented DMEM prior to implantation. The average plug size was 1.98 ± 0.05 mm in diameter and 3.10 ± 0.12 mm in length (*N* = 12). Sham plugs were identically prepared using A3G7 hydrogel without cancer cells.

#### Animals and surgical procedures

Ten male SRG™ OncoRats (Charles River Laboratories), aged 10 weeks at the time of surgery, with one rat dying prematurely due to megaesophagus. Rats were housed in pairs under a 12-hour light/dark cycle with ad libitum access to water and a wet mash diet to reduce the risk of megaesophagus, a strain-associated complication. All procedures were approved by the Research Institute of the McGill University Health Centre (RI-MUHC) Animal Care Committee (protocol #MUHC-8095).

All surgical procedures were conducted under aseptic conditions within a Class II biosafety cabinet (BSC). Preoperative analgesia consisted of sustained-release buprenorphine (1 mg/kg, SC) and carprofen (5 mg/kg, SC), administered 30 minutes before anesthesia induction. Anesthesia was maintained using isoflurane (2-2.5%) delivered in oxygen via nose cone. Anesthetic depth was monitored by pedal withdrawal reflex, and core temperature was maintained using a feedback-controlled heating platform (PhysioSuite, Kent Scientific). All rats received subcutaneous warmed saline (0.2 mL/10 g) after surgery and were monitored twice daily for 48 hours. Body weights were recorded every 2-10 days. Sutures were removed 7-10 days postoperatively. Representative intraoperative images of Surgery 1 and Surgery 2 are shown in Figure 3.

**Figure 3.**
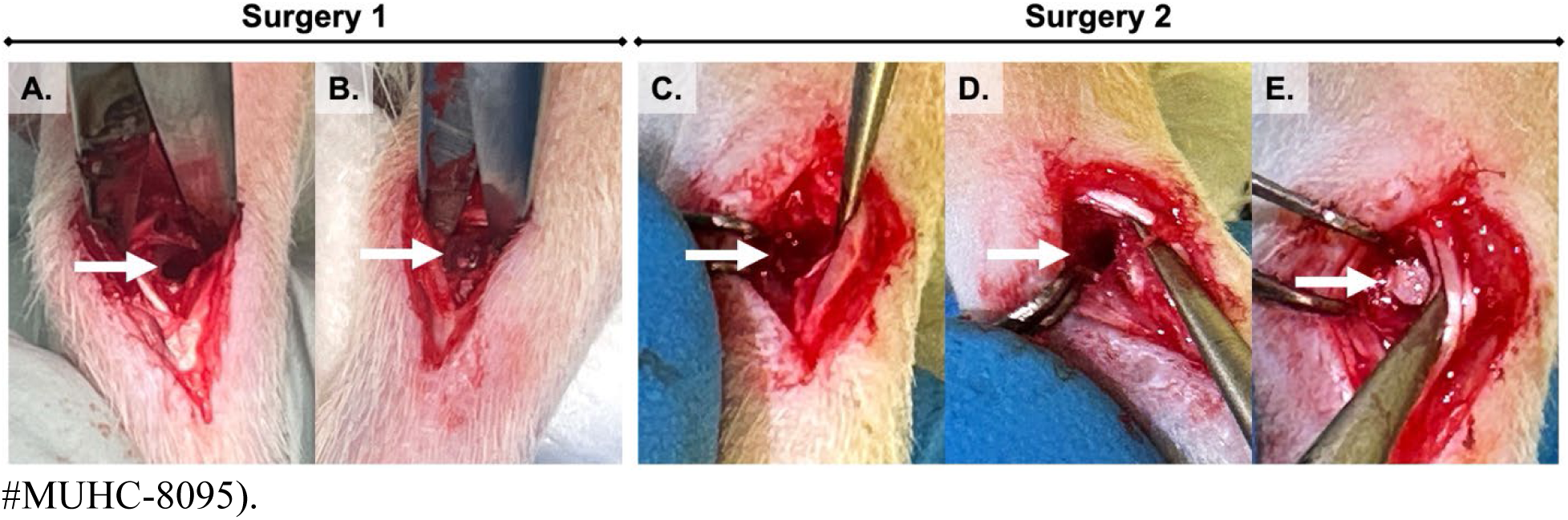
Surgical procedure for tumor implantation and cement treatment in the rat caudal vertebral model. **A**. A 1.8 mm hole is drilled into the caudal vertebral body (arrow) during Surgery 1; **B**. Tumor plug is inserted into the drilled cavity (arrow); **C**. Six weeks post-implantation, the tumor plug is visibly integrated at the vertebral site (arrow), prior to resection (Surgery 2); **D**. Following tumor resection, the vertebral defect is slightly widened using a surgical burr to ∼2.8-3mm maximum (arrow); **E**. Implant is press-fit within the prepared cavity (arrow) to complete the treatment procedure.

#### Surgery 1: Tumor plug implantation

Nine male SRG™ OncoRats (Charles River Laboratories), aged 10 weeks, underwent tumor implantation at the proximal caudal vertebra. Animals were positioned prone. Animals were positioned prone. The target vertebral level was localized by manual palpation of the caudal spine, identifying the intervertebral disc spaces by their increased flexibility between adjacent vertebrae. A 1 cm dorsolateral skin incision was made approximately 1 cm distal to the tail base. Minor bleeding was controlled with sterile gauze. Blunt dissection with fine-tip forceps was used to expose the paraspinal musculature and the vertebral body. Using a high-speed micro-drill (Ideal Micro-Drill™, CellPoint Scientific) fitted with a 1.8 mm carbide ball-tip burr, a 2.5-3.0 mm deep cylindrical intraosseous cavity was created into the lateral vertebral cortex. Tumor plugs were inserted flush with the surrounding cortical surface using forceps. Two additional rats received sham implants consisting of blank A3G7 hydrogel plugs (no cells), inserted into identical drill defects following the same protocol. These served as negative controls for tumor growth, (not shown). The muscle and tendon layers were re-apposed, and the site was flushed with sterile saline and locally infiltrated with lidocaine/bupivacaine. The skin was closed with interrupted 4-0 Monocryl suture (Ethicon). For bioluminescence imaging (BLI), 150 mg/kg of body weight of D-luciferin (Thermo-Fischer, ON Canada – prepared with sterile filtration in PBS) was injected intraperitoneally with a 25-gauge 1cc sterile needle/syringe, under sedation. After 20 minutes of incubation time, rats were imaged using a Bruker In-vivo Xtreme (Bruker, Burlington ON, Canada).

#### Surgery 2: Tumor resection and cement implantation

Three weeks after implantation and BLI inspection, all rats underwent tumor resection in the same BSC environment. The previous incision site was reopened, and soft tissues were carefully re-dissected to access the tumor mass. The tumor was debulked using intralesional curettage with fine-tip forceps. A microdrill was used to slightly expand the bone defect about 0.5mm. Cylindrical composite 3D printed scaffold implants were disinfected by overnight soaking in 70% ethanol, followed by 3 washes in sterile PBS. Scaffolds were then exposed to 15 minutes of UV treatment for each side and then dried in the biosafety cabinet. Half of the scaffolds were loaded in the LayFomm central part with 500 μM doxorubicin and dried for an additional 1 hour. Scaffolds were inserted press-fit as best possible into the bone defect according to group allocation: control scaffold (*n* = 4) or DOX-loaded scaffold (*n* = 5). The surgical field was again irrigated, and closure was performed. All rats were euthanized 6 weeks post-implantation by isoflurane followed by CO₂ overdose and pneumothorax. Tails were then harvested and fixed in 10% neutral buffered formalin for micro-CT analysis.

### Statistical Analyses

Statistical tests were performed using GraphPad Prism 9.0 (GraphPad Software, La Jolla, CA, USA). Each experiment was performed thrice and in triplicate. The results are reported as mean ± SEM with an n = 4 control and 5 dox for animal studies, and n=3 for all other experiments unless noted otherwise. Student’s t tests were performed to test for significance between the mechanical properties of Lactoprene and composite scaffolds, as well as between the total amounts of doxorubicin released from scaffolds loaded with 175 and 350ng of doxorubicin. One-way ANOVAs were performed to test for significance between drug-loaded scaffold treatments. Dose-response data were fitted with sigmoidal functions and the IC_50_ values were determined by Prism. Statistical significance was taken as *p* < 0.05. The significance reported are the following: not significant (ns) *p* > 0.05; * *p* < 0.05; ** *p* < 0.01; *** *p* < 0.001; **** *p* < 0.0001.

## Results

### Scaffold Fabrication and Compression Testing

The prepared composite Lay FOMM/Lactoprene scaffolds were approximately 2 mm in height and 3 mm in diameter. To assess the mechanical properties of the scaffolds, compression testing of both the outer Lactoprene shell (*n* = 5) and the composite scaffold (*n* = 9) was performed. Stress-strain curves were plotted (Figure 4a), and the mechanical properties were calculated. The Young’s moduli of the Lactoprene shell and composite scaffolds were 198.96 MPa (±48.49 MPa) and 154.57 MPa (±58.86 MPa), respectively, however there were no statistically significant differences (Figure 4b, *p* > 0.05). The yield stress was 8.36 MPa (±0.88 MPa) for Lactoprene and 7.41 MPa (±2.85 MPa) for composite scaffolds. Similarly, there were no significant differences between the yield stresses (Figure 4c, *p* > 0.05). The composite scaffolds demonstrated an increase in the yield strain, which was calculated as a percent yield strain, from 5.56% (±0.94%) for Lactoprene scaffolds to 8.77% (±0.83%) for composite scaffolds, which was found to be a statistically significant difference in the amount of compression to reach the yield point (Figure 4d, *p* < 0.0001).

**Figure 4.**
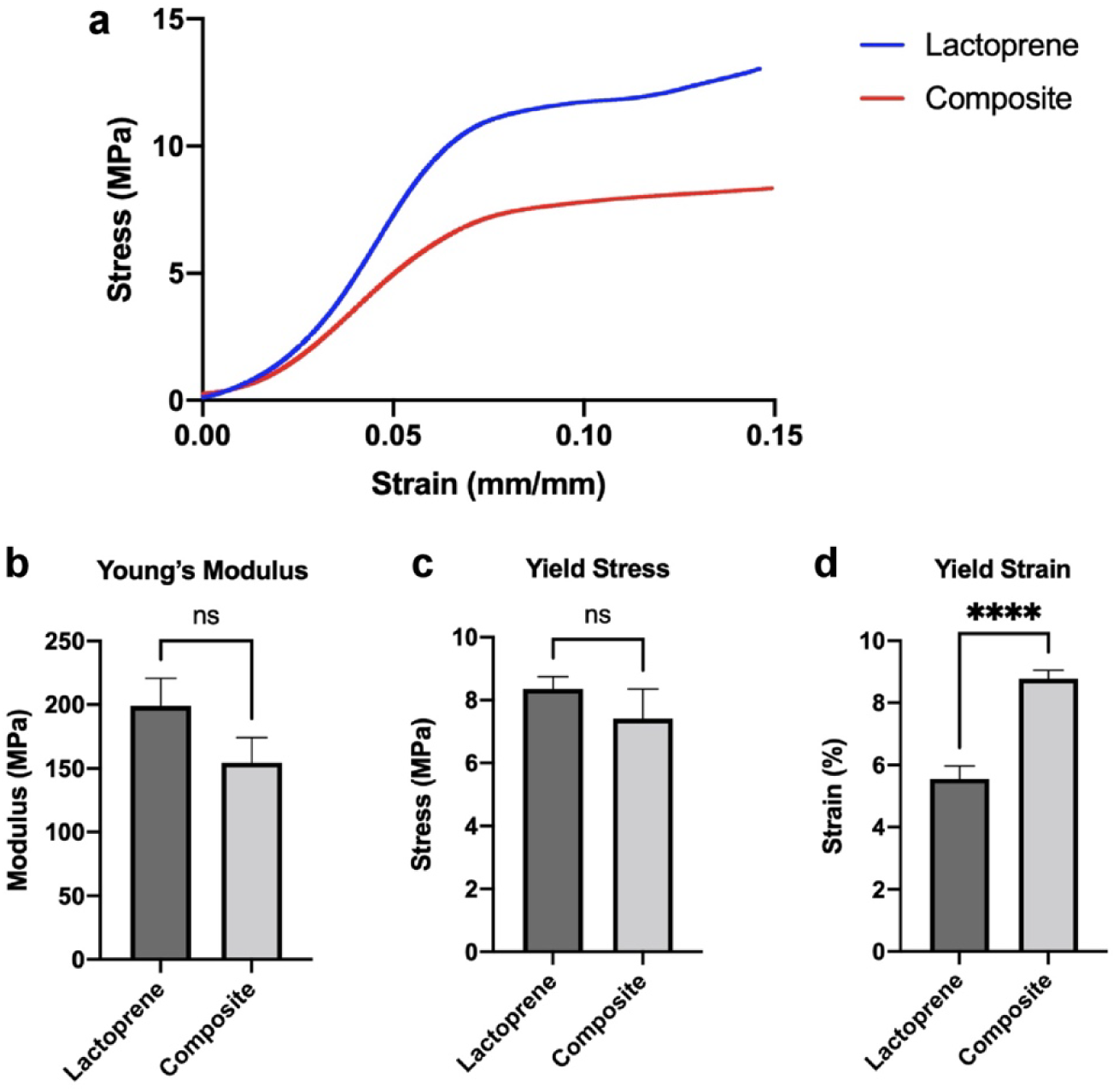
Compression testing was done on Lactoprene and composite scaffolds, yielding (a) stress-strain curves for Lactoprene (*n* = 5) and Composite scaffolds (*n* = 9). The three mechanical properties displayed are: (b) Young’s modulus, (c) yield stress, and (d) yield strain.

### Cumulative Doxorubicin Release from 3D Printed Scaffolds

We previously measured doxorubicin release from small scaffolds made of Lay FOMM 60 compared to other 3D printed scaffolds [20]. To determine the release kinetics when the Lay FOMM is part of a composite structure, we performed a similar drug release experiment. The release rate of doxorubicin from 3D printed composite scaffolds was quantified over a period of 7 days (Figure 5). The proportional release was not significantly different at the end of the 7 days between scaffolds loaded with 175 ng and 350 ng of doxorubicin (Figure 5a, *p* > 0.05). The initial release of the scaffolds loaded with a higher amount of drug was higher, however, the difference became insignificant over the course of 7 days. Scaffolds loaded with 175 ng released 100.02 ng (±16.55 ng), which was 57.2% of the loaded amount, and those loaded with 350 ng released 226.53 ng (±19.98 ng), which was 64.7% of the loaded amount, in 7 days (Figure 5b, *p* < 0.0001). Therefore, with more doxorubicin loaded, there is a significantly higher amount of doxorubicin released. However, the proportion of doxorubicin released did not significantly differ between the loading doses.

**Figure 5.**
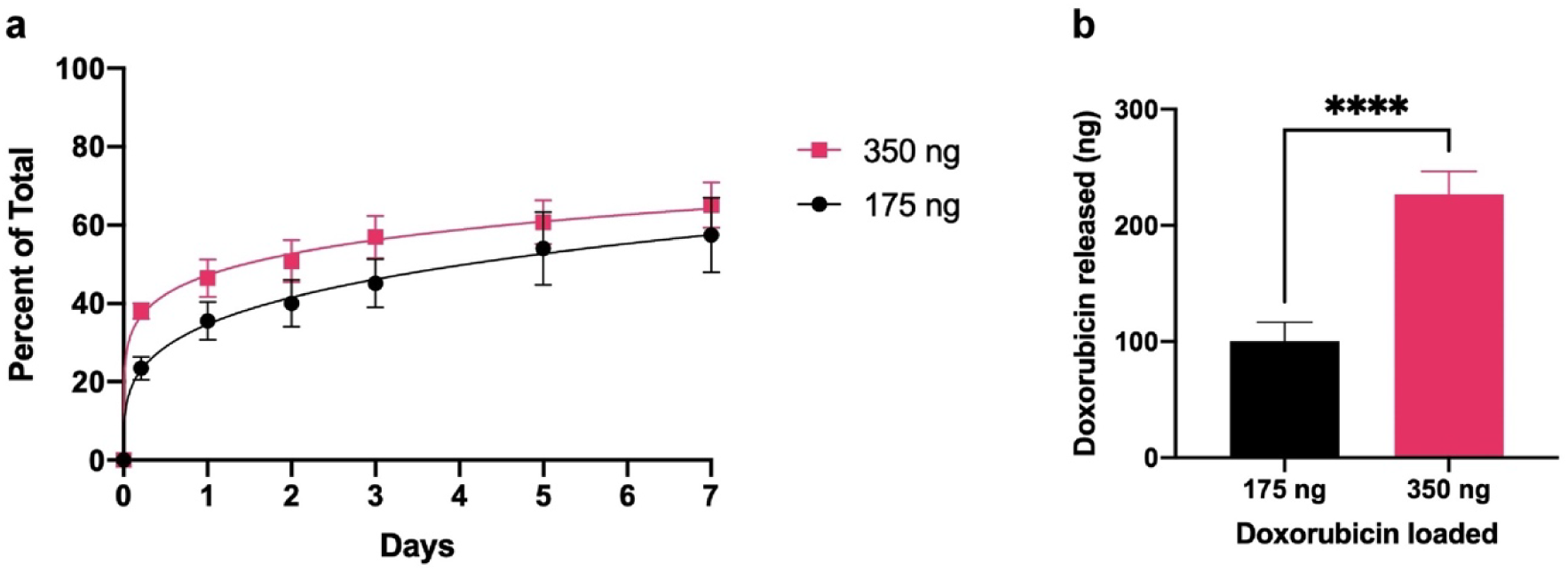
(a) Cumulative doxorubicin release from 3D printed composite scaffolds loaded with differing amounts (175, 350 ng) over the course of 7 days, expressed in percent of total drug loaded. (b) Amount of doxorubicin loaded as a function of doxorubicin released, expressed in nanograms. Data reported as mean ± SD with *n* = 4.

### Dose-Response Curves

To determine the optimal drug loading concentrations for the 3D printed scaffolds, dose-response assays were conducted on MDA-MB-231/GFP and C4-2B cell lines. The MDA-MB-231/GFP cell line displayed a higher tolerance against both doxorubicin and cisplatin compared to the C4-2B line, with the relative IC_50_ values being 1.076 µM for doxorubicin and 34.21 µM for cisplatin (Figure 6a-b). The relative IC_50_ values for C4-2B cells for doxorubicin and cisplatin were determined to be 0.2651 µM and 7.933 µM, respectively (Figure 6c-d). Based on the drug release profile and IC_50_ values obtained, the 3D printed scaffolds were loaded with doxorubicin and cisplatin such that the released concentrations would fall in the therapeutic range. These loading concentrations are outlined in Table 1 and Table 2.

**Figure 6.**
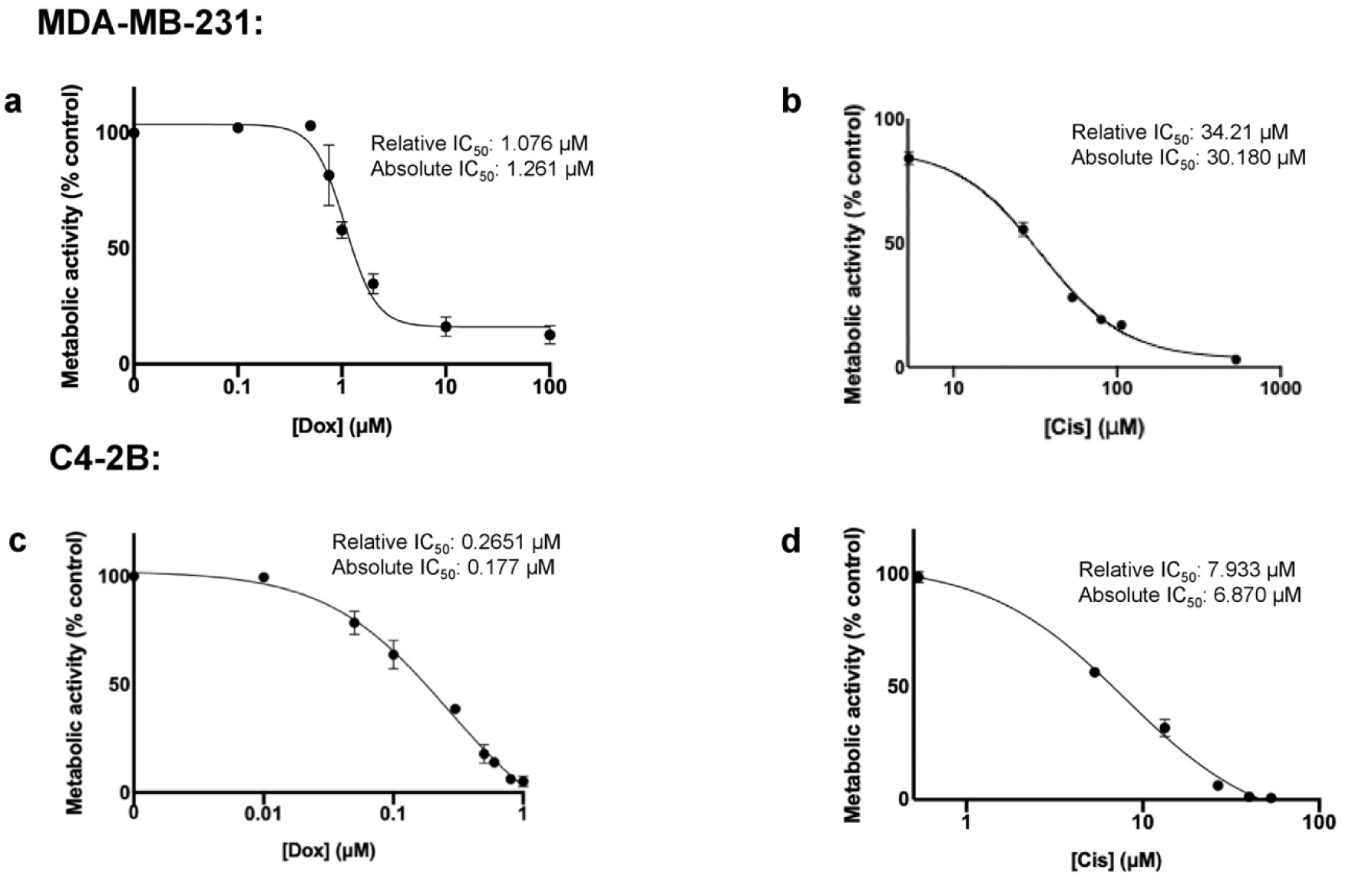
Relative IC_50_ curves determined for MDA-MB-231/GFP cells treated with chemotherapeutics (a) doxorubicin, and (b) cisplatin, as well as for C4-2B cells treated with (c) doxorubicin and (d) cisplatin.

### 3D Scaffold Drug Delivery to MDA-MB-231/Luc and C4-2B Cells

The drug-loaded scaffolds were incubated with the two cell lines to assess *in vitro* treatment efficacy. The doxorubicin release rate data were used to determine for dosing the scaffolds. For cisplatin, we used several cisplatin concentrations to create a standard curve in order to determine the optimal cisplatin release to kill at least 50% of cells. MDA-MB-231/Luc cells treated with doxorubicin-infused composite scaffolds for 5 days showed a 87.64% (±2.52%) reduction in cell metabolic activity compared to cells treated with the control scaffolds loaded with PBS (Figure 7a, *p* < 0.0001). Similarly, all cisplatin-loaded scaffolds significantly inhibited MDA-MB-231/Luc cell activity, indicating more than 50% cell death, with a 63.77% (±2.57%) to 97.8% (±0.18%) reduction in activity depending on the loaded dose (Figure 7b, *p* < 0.0001). C4-2B cells treated with doxorubicin loaded scaffolds showed a 72.03% (±11.03%) reduction in metabolic activity compared to the control condition (Figure 7c, *p* < 0.0001). Likewise, all cisplatin-infused scaffolds significantly inhibited C4-2B cell activity by 72.82% (±3.82%) to 97.35% (±1.75%) (Figure 7d, *p* < 0.0001). Overall, higher loading doses of cisplatin provided a stronger effect with regard to cell metabolic activity inhibition. The metabolic activity of positive controls for both direct doxorubicin and cisplatin treatments were comparable to drug-infused scaffold treatments for both MDA-MB-231/Luc and C4-2B cell lines.

**Figure 7.**
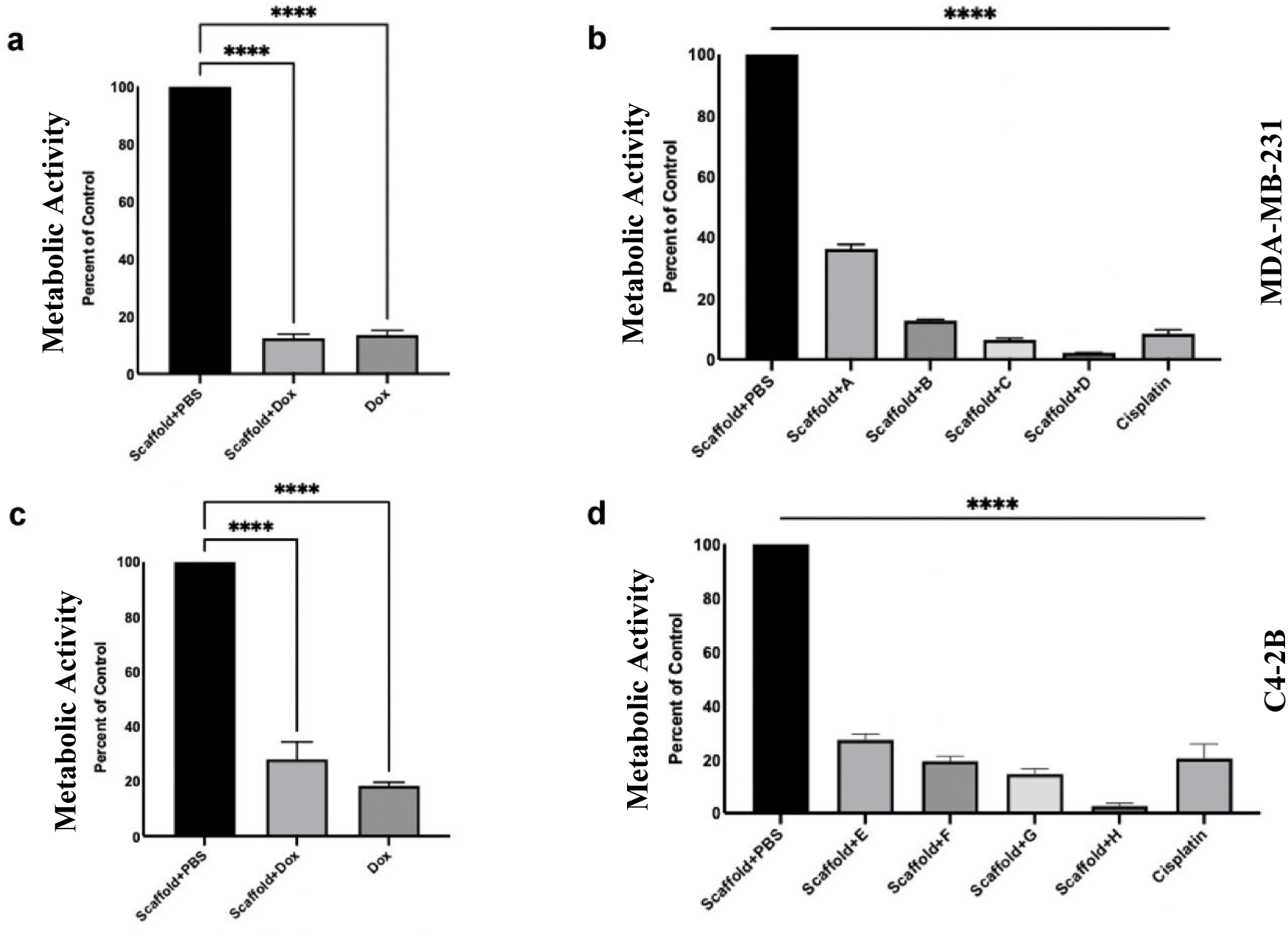
Metabolic activity of breast and prostate cancer cells measured with AlamarBlue assay after 5 days of treatment with 3D printed composite scaffolds loaded with PBS, doxorubicin(dox) or cisplatin and direct drug treatment, expressed in percent of control. (a) MDA-MB-231/Luc cells with concentrations of 1 µM of free dox and 150 µM of dox-infused composite scaffolds; (b) MDA-MB-231/Luc cells with concentrations of 32 µM of free cisplatin and varying cisplatin doses loaded onto composite scaffolds; (c) C4-2B prostate cancer cell line with concentrations of 0.2 µM of free dox and 30 µM of dox-infused composite scaffolds; (d) C4-2B cells with concentrations of 8 µM of free cisplatin and varying cisplatin doses loaded onto composite scaffolds. Refer to Tables 1 and 2 for scaffold cisplatin concentrations.

### 3D Scaffold Drug Delivery in Mixed Migration Model

We previously developed a mixed migration 3D model to assess efficacy of drug delivery against cancer cell migration to a tissue interface [22]. Figure 8 briefly describes the experimental design and evaluation. To assess efficacy against cell activity and cell migration of the drug-loaded scaffolds in a physiologically-relevant 3D model, scaffolds were placed in our previously described PP-3D-S co-culture model with MDA-MB-231/GFP tumor cells and IMR-90/mCherry fibroblasts. When treated with doxorubicin-loaded scaffolds, fluorescence microscope images revealed a lower number of migrated tumor cells compared to direct doxorubicin treatment controls (Figure 9A). The reduction of cell metabolic activity with doxorubicin-loaded composite scaffolds and direct doxorubicin treatments (positive controls) was similar. The metabolic activity of the breast cancer cells was assessed in the PP-3D-S models with drug-infused 3D printed composite scaffolds and direct media supplementation treatment. The controls that were used as comparison for the 3D printed composite scaffolds were 3D printed scaffolds loaded with PBS, and for the direct treatment groups the controls were the PP-3D-S scaffolds in media with no media supplementation. Both 3D printed scaffolds and direct treatment significantly inhibited metabolic activity at all doses tested compared to their respective controls (Figure 9b, *p* < 0.0001). Finally, the number of migrated tumor cells were counted and demonstrated a 60.80% (±4.10%) to 66.69% (±1.42%) reduction in cell migration with doxorubicin-infused 3D printed composite scaffold treatment compared to controls (Figure 9c, *p* < 0.01).

**Figure 8.**
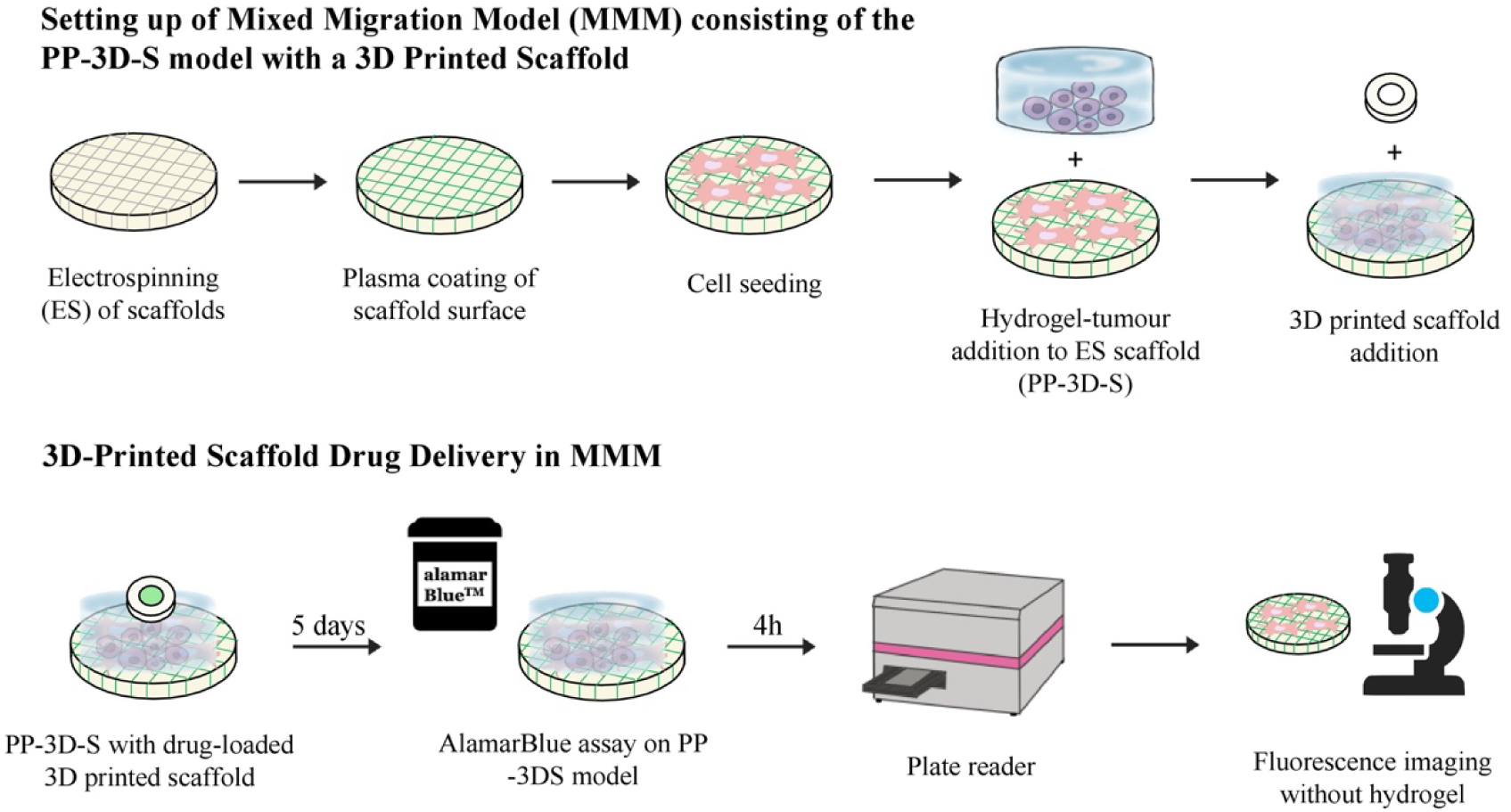
Experimental design using PP-3D-S scaffolds with 3D printed drug-infused scaffolds for in vitro treatment of breast cancer cells, assessing metabolic activity via plate reader and migration via fluorescence imaging.

**Figure 9.**
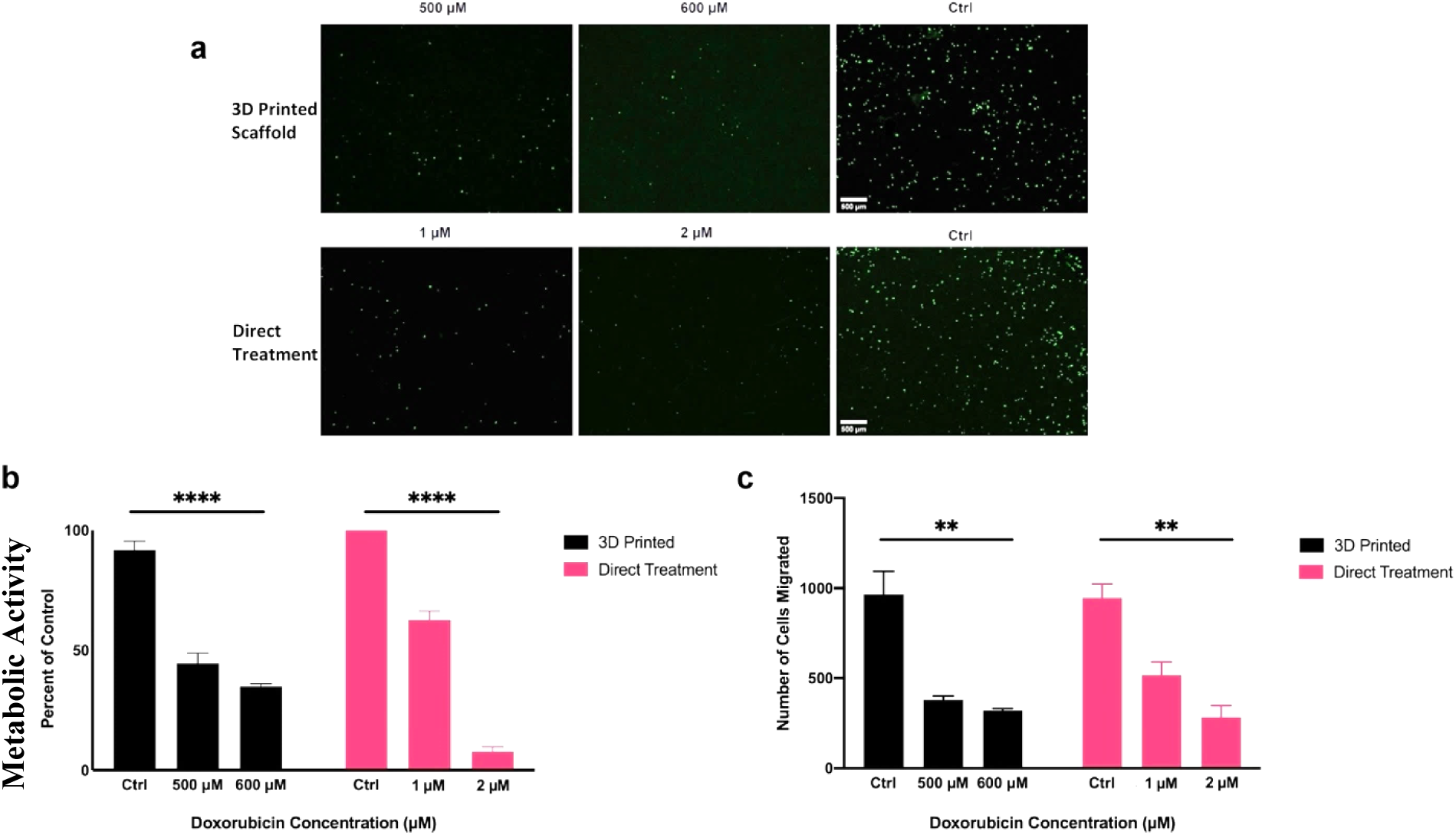
3D printed composite scaffold drug delivery in Mixed Migration Model (MMM). (a) Fluorescence microscope images of migrated breast cancer cells with 3D printed scaffolds loaded with doxorubicin at 500 µM (*n* = 3) and 600 µM (*n* = 2)(based on release kinetics from our previous study [16]), and direct treatment of 1 µM (*n* = 3) and 2 µM (*n* = 3) of doxorubicin. The controls were a scaffold loaded with PBS for the 3D printed condition and media with no drug for the direct treatment. (b) Metabolic activity of MDA-MB-231/GFP cells measured with AlamarBlue assay after 5 days of treatment with 3D printed and direct treatment at previously mentioned doses of doxorubicin compared to respective controls, expressed in percent of direct treatment control. (c) Number of migrated tumour cells counted with the same experimental and control groups and doses previously mentioned.

### In Vivo Feasibility Assessment of Implanted Scaffolds

To test the feasibility of using the composite 3D printed scaffold for local drug delivery and tissue regeneration in tumor-resected bone defects, we established an *in vivo* mimic of spine metastasis. The pipeline for this *in vivo* approach is outlined in Figure 10. Following implantation of bioluminescent human triple negative breast cancer cells into caudal vertebrae of SRG rats, tumor growth could be observed over 3 weeks using live-imaging (Figure 11A). Following tumor resection and widening of the surgical resection bed by about 0.5mm, the composite 3D printed scaffolds were embedded and after 6 weeks of implantation, bone integration was assessed. MicroCT analysis revealed that both drug-loaded and control scaffolds had similar bone volume per tissue volume at the interface between the scaffolds and vertebral bone, indicating no observable negative impact of scaffolds (Figure 11B). Live imaging prior to euthanasia did not reveal any observable signal for tumor cells in controls or drug treated animals (data not shown), as six weeks of follow was seemingly insufficient for enough growth. Together, these data indicate that implantation of the 3D printed composite scaffolds is feasible and does not cause adverse events over the time-course of the experiment.

**Figure 10.**
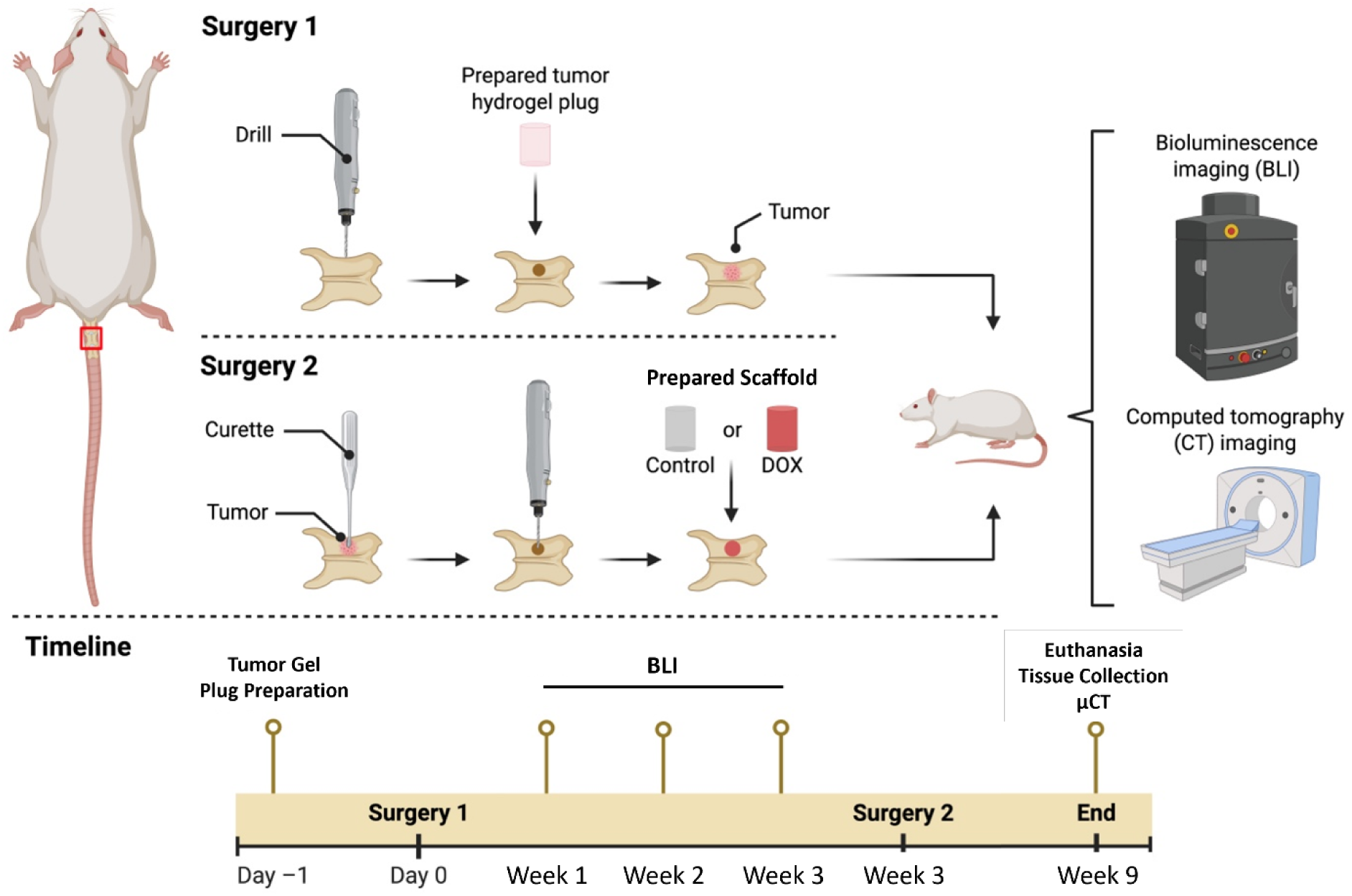
Schematic showing the in vivo experimental workflow. Surgery 1 is performed and tracked for 3 weeks, at which point surgery 2 occurs. Control and test scaffolds are implanted, animals are followed for 6 weeks, and then vertebrae are recovered for microCT analysis.

**Figure 11.**
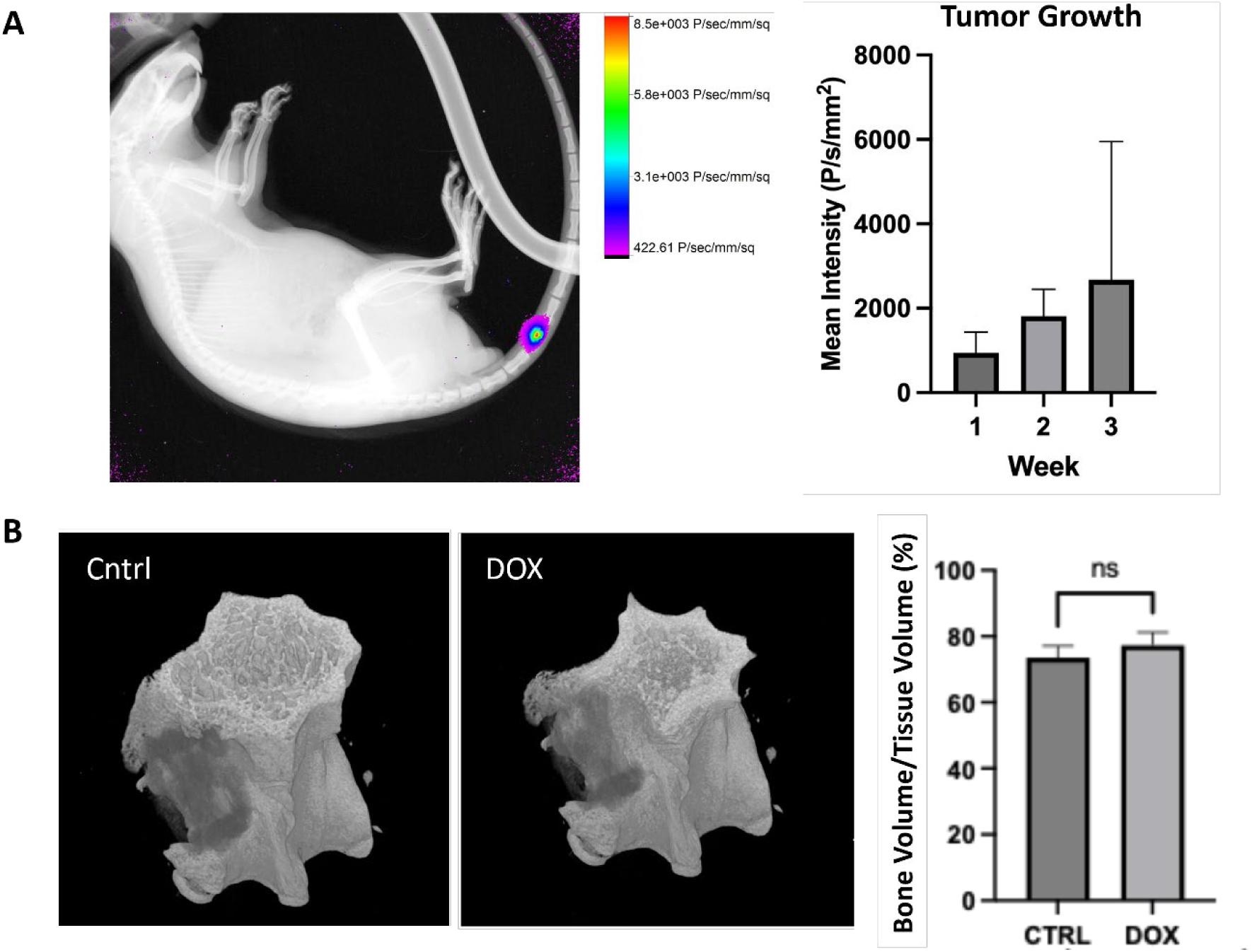
In vivo feasibility testing of drug-loaded composite 3D printed scaffolds. A) representative live-bioluminescence image of triple-negative breast cancer cells 3 weeks post-implantation into the caudal vertebrae representing a spine metastasis. Growth is observed over 3 weeks as indicated in graph. B) Representative 3D reconstructed microCT images showing implants on left side of vertebrae. Bone volume/tissue volume quantification at the interface of scaffolds and bone is shown in the graph, indicating no difference. Error bars = ± SD. n= 4 control and 5 test animals.

## Discussion

The purpose of this study was to determine whether 3D printed composite scaffolds have comparable mechanical strength to trabecular bone and whether composite scaffolds loaded with chemotherapeutics will be equally effective at locally inhibiting tumor cell proliferation and migration *in vitro* compared to direct treatment (simulating systemic delivery). To assess this, compression testing of composite scaffolds, drug release measurement, IC_50_ determination, and 2D and 3D treatment with composite scaffolds were conducted. We show feasibility of drug loading for two standard chemotherapuetics, release and efficacy against one breast cancer and one prostate cancer cell line.

Mechanical testing yielded no significant difference in Young’s modulus between the Lactoprene and composite scaffolds (*p* > 0.05). Given this, we can conclude that the addition of Lay FOMM in the composite scaffolds did not significantly alter the stiffness of the material, despite Lay FOMM’s flexibility. We also found no significant differences on yield point or stress between Lactoprene and Lay FOMM, indicating that the addition of Lay FOMM does not significantly change the yield stress of the scaffolds (*p* > 0.05). On the other hand, we found that the percent strain at yield was significantly higher for the composite scaffolds compared to Lactoprene alone (*p* < 0.0001), attributable to LayFOMM’s porous, sponge-like nature. The composite scaffolds are higher in elasticity compared to Lactoprene-only scaffolds, which may provide a cushioning effect in the bone to stabilize it without the addition of isolated pressure points.

Comparing the mechanics of our developed scaffolds to trabecular bone is not trivial, namely due to variations in bone volume fraction, individual arrangement of trabeculae, aging, and disease [25]. Due to this, there is a wide range of mechanical properties of trabecular bone found in the literature . One previous study determined a Young’s modulus of vertebral trabecular bone of 344 MPa (±148) [26]. Other studies have found the Young’s modulus to be as low as 67 MPa (±45) [27]. In the current study, the Young’s modulus was determined to be 198.96 MPa (±48.49) for Lactoprene and 154.57 MPa (±58.86) for composite scaffolds, which are within the range of what was found in the literature for vertebral trabecular bone. In one study, the yield stress was determined to be 2.02 MPa (±0.92), and the yield strain to be 0.77% (±0.06) [26]. The stress is higher in the scaffolds at the limit of elastic behaviour compared to trabecular bone, indicating that they have the necessary capacity to bear weight.

Composite scaffolds loaded with doxorubicin released approximately 60% of the loaded doxorubicin within a week, regardless of the amount loaded. The proportional release was nearly identical even with double the amount of loaded doxorubicin, indicating that the release is heavily dependent on the properties of the material used for drug delivery. Previous work in our group has indeed demonstrated that the drug release kinetics can be modified with the selection of different materials, with more porous materials leading to a higher proportional release over a period of one week [16, 17]. Our results demonstrated that the release of cisplatin from the scaffolds was effective at inhibiting the MDA-MB-231 and C4-2B cells *in vitro*, and that the effect was comparable to that of the direct cisplatin treatment. Therefore, the composite Lactoprene/Lay FOMM scaffolds were effective at least for sequestering and releasing two chemotherapuetics *in vitro*. Dang *et al*. fabricated 3D printed scaffolds with micro- and macro-scale pores using polycaprolactone (PCL) mixed with porogen microparticles [28]. They determined that the microscale porosity decreased the rapid initial release (burst release) of doxorubicin [28]. Multiple groups have 3D printed calcium phosphate cements for drug delivery applications. Wu *et al*. explored calcium phosphate for the delivery of 5-fluorouracil, another anti-cancer drug, and the treated scaffolds released 100% of the loaded drug within 2 hours [29]. Drug release from calcium phosphate cement occurs rapidly in a burst release manner and is typically limited to a total release lasting less than 24 hours [30]. In either case, this would not be conducive to treating spine metastases resections, since disseminated tumor cells can remain dormant in the bone marrow niche only to recur sometime later. A sustained release device, like the one presented in the present study, would be more appropriate for such treatments. Zhu *et al*. combined mesoporous bioactive glasses and mesoporous silica nanoparticles with poly (3-hydroxybutyrate-co-3-hydroxyhexanoate) in a composite 3D printed scaffold which was mesoporous and sustainably released anti-tuberculosis drugs compared to calcium phosphate cements [31]. Sustained release is more favorable in comparison to burst release as it allows for long-term treatment and can suppress tumor growth at higher doses. In addition, burst release might be toxic for the adjacent healthy cells. In combination with our results, this demonstrates the importance of porous structures for 3D printed scaffolds intended for prolonged drug delivery applications and decreasing the burst release.

The IC_50_ values were determined to be 1.076 µM and 34.21 µM for MDA-MB-231 cells treated with doxorubicin and cisplatin, respectively, and 0.2651 µM and 7.933 µM for C4-2B cells treated with doxorubicin and cisplatin, respectively. Our determined values were similar to the range of values found in past studies on MDA-MB-231 cells treated with doxorubicin between 3 - 23 µM [32–35] and those treated with cisplatin, ranging from 3 to 30 µM [36, 37]. Some discrepancies in dose results may be attributed to using MTT versus Alamar metabolic assays. Our findings on C4-2B sensitivity to doxorubicin were close to what has been reported for IC_50_ in the literature - 32 nM [38]. Overall, the IC_50_ values determined in this paper were similar to what was found in the literature for MDA-MB-231 and C4-2B cells, indicating confidence in our subsequent drug release experiments.

Once chemotherapeutic IC_50_ doses were established for the MDA-MB-231 and C4-2B cells, composite scaffolds were loaded with doxorubicin and cisplatin separately for release assessment. The impact on metabolic activity for both doxorubicin and cisplatin release scaffolds was significantly lower compared to untreated controls and very comparable to positive controls (direct chemotherapeutic treatments) in both cell lines. This demonstrated appropriate therapeutic release and delivery at this minimal *in vitro* level. The novelty in the present study is the use of Lay FOMM for uptake and subsequent release of the drugs within a composite 3D printed material. Furthermore, our developed construct has a sustained release profile, effective for inhibiting the metabolic activity of multiple cancer cell types. A similar study demonstrated that 3D printed calcium phosphate scaffolds can be loaded with curcumin-liposomes in bone repair applications in osteosarcoma models. The release of curcumin from the scaffolds promoted human fetal osteoblasts viability while inducing death to human osteosarcoma cells [39]. Another study showed that 3D printed gelatin-based implant could release chemotherapeutics and growth factors for osteogenesis and anti-tumor therapy, which inhibited breast cancer and osteosarcoma tumor growth and promoted bone stability in vivo [40]. Along with the current study, these other studies support the notion that 3D printed scaffolds can be loaded with therapeutics to fill bone defects, effectively release anticancer therapeutics, and promote bone health.

To assess the impact of drug-loaded 3D printed scaffolds on metastatic events, we applied our drug-loaded scaffolds to an established 3D co-culture bone metastasis model [22–24]. This type of 3D model is more physiological and allows better cellular responses to drugs eluted from the 3D composite scaffolds [41]. We determined doxorubicin release from the composite 3D printed scaffolds was equally effective to direct treatment on blocking MDA-MB-231 cell migration from the hydrogel compartment to the bone-like compartment. This also correlated with comparable reductions in metabolic activity following treatments. The inhibition of metabolic activity and cell migration was dose-dependent, with higher doses leading to a stronger inhibitory effect. This demonstrates the ability of the scaffolds’ drug release to impair cell activity and motility. To our knowledge, studies testing the effect of drug-loaded 3D printed scaffolds on cells in 3D culture models are limited. Zhang *et al*. prepared composite PCL/hydrogel 3D printed scaffolds loaded with resveratrol and strontium ranelate for wound healing assays. [42]. Interestingly, scaffolds loaded with resveratrol increased vascular endothelial cell migration driving higher wound closure after 16h. However, the release could be carried out for 21 days with 30% of the loaded molecule released [42]. Our scaffolds could deliver more drug over this same time frame, with room for potentially more control using different polymer variations [16]. Other studies have shown that drug loaded 3D printed scaffolds can enhance mesenchymal stromal cell migration in wound healing assays as well [43]. These studies point out that our scaffolds could benefit from loading additional drugs like growth factors, which could support bone health and regeneration in addition to blocking cancer invasion and growth in the bone. Studies assessing the efficacy of chemotherapeutic-loaded 3D printed scaffolds on cancer cell migration are limited, and typically involve the use of Boyden chamber devices, which lack certain physiological aspects. Future *in vivo* studies would best be suited for evaluating cell motility/metastases within the bone and are required to predict the success of these drug delivery devices.

We showed in this study the feasibility of using the lactoprene-βTCP-LayFomm compositive scaffolds *in vivo* to fill a bony defect in a novel rat model of spine metastasis. Both the lactoprene and Lay-Fomm materials have been shown to be compatible in vivo in our previous work [20, 44] as well as some various in vitro tests by others [45–47]. The use of such polymers offers several advantages, including tunable mechanical strength, controlled porosity as small as 0.5mm (using conventional desktop 3D printers), and customizable degradation profiles that can be optimized to match the physiological requirements of bone healing. Our previous studies have also indicated the cytocompatibility and biocompatibility of the individual polymer components. Incorporating drug-loaded components such as the LayFomm, into the lactoprene structure gives the additional advantage for sustained and local drug release of anti-cancer therapies while minimizing toxicity associated with systemic delivery. It would be of value to test the release of osteogenic, antimicrobial, or anti-inflammatory therapies in the future as well. While we focused on testing the feasibility of using the composite scaffolds *in vivo* in a pilot test here, it remains necessary to evaluate print fidelity, structural stability, drug loading efficiency, release kinetics, long-term biocompatibility, and the scaffold’s ability to support cellular attachment and bone tissue formation in a higher-powered future study. Further, established a metastatic spine recurrence timeline will also be of value in future work. At the very least, our small cohort study has established several novel *in vivo* parameters which provide essential proof-of-concept data supporting the translational potential of our composite 3D printed scaffolds.

A key limitation of this research lies in the use of a commercial material of which the precise composition is unknown. The 3D printed scaffolds were loaded with doxorubicin and cisplatin on a single side, which may not have uniformly dispersed the drug solution throughout the Lay FOMM core. The doxorubicin release kinetics were evaluated using a fluorescence microplate reader and interpolated from a standard curve, with samples taken at specified time points. Varying the volume of the aliquot taken and the volume replenished in the samples may affect the release rates of the drug, as the sink conditions may be affected and produce variable readout results. Future studies may assess drug release with varying sink conditions, as well as various release medium other than PBS. Cisplatin release was measured indirectly by measuring cell metabolic activity in response to the loaded scaffolds as the drug is not fluorescent. Subsequent studies may derivatize cisplatin with o-phenylenediamine to quantify it spectrophotometrically [48]. Furthermore, prolonging the experimental duration for measuring drug release kinetics would allow for a more complete release profile. The 2D and 3D *in vitro* cell metabolic activity and cell migration assays relied on the use of cell lines, which may not accurately depict tumor cells in their native tumor microenvironments. Therefore, use of patient-derived cells would strengthen the results and upon comparison to patient treatment data, may indicate whether our devices are appropriate for personalized medicine. Additionally, there is a possibility that the Lactoprene shell would not integrate with the bone in an *in vivo* model. It has not been established whether the chemotherapeutics would damage the surrounding healthy tissue, thereby preventing osseointegration. However, the chemotherapeutics specifically target rapidly dividing cells, making it less likely that osteoblasts and other bone cells would be targeted, and using a Lactoprene-β-TCP composite in an *in vivo* model could promote bony integration, as in our previous study [44]. Finally, the scaffolds’ ability to attach cells and integrate into the target tissue, as well as the degradation rate of the scaffolds may be evaluated to assess osseointegration and safety *in vivo*. Our earlier studies have indicated that Lactoprene can readily bind MSCs [49] and osteoblasts, and that Lay FOMM can be bio-resorbed [20].

With the recent rise in cancer incidence and increased risk of recurrence and reoperation of metastatic bone lesions, the current surgical gold standard may be improved with 3D printed scaffolds for bone repair and drug delivery. We demonstrated here that 3D printed scaffolds can be manufactured at a low cost with mechanical properties similar to trabecular bone. Furthermore, the 3D printed composite scaffolds were loaded with doxorubicin and cisplatin chemotherapeutics and effective doses were released against breast and prostate cancer cells lines, inhibiting cell activity and migration. An *in vivo* tumor metastasis model will allow for further pre-clinical testing of this drug delivery device to assess efficacy and ability to regenerate bone.

## Acknowledgments

This project was funded by the Réseau de recherche en santé buccodentaire et osseuse (RBSO) Major Structuring Young Investigator Grant awarded to MHW, IV, LH and DHR. We are thankful for support from Transplant Quebec for patient specimens. DHR gives special thanks to continued support from the McGill Scoliosis and Spine Group. AP and AS both received studentships from the Research Institute of McGill University Health Centre, FRQS and CIHR. MMG received fellowships from the Research Institute of McGill University Health Centre and the Montreal General Hospital Foundation/CODE LIFE.

## Author Contributions

conceptualization – AA, MW, MHW, IV, LH and DHR; data curation – AAP, AS, and MMG; investigation – AAP, AS and MMG; formal analysis – AAP, AS, MMG and BNB; supervision – IV and DHR; funding acquisition – MHW, LH, IVand DHR; methodology – AAP, AS, MMG, AA and MW; writing original draft – AAP, AS and BNB; writing editing and review – all authors.

## Data

The data generated in this study are available upon request from the corresponding authors.

